# Cross-β helical filaments of Tau and TMEM106B in Gray and White Matter of Multiple System Tauopathy with presenile Dementia

**DOI:** 10.1101/2023.01.09.523314

**Authors:** Md Rejaul Hoq, Sakshibeedu R. Bharath, Grace I. Hallinan, Anllely Fernandez, Frank S. Vago, Kadir A. Ozcan, Daoyi Li, Holly J. Garringer, Ruben Vidal, Bernardino Ghetti, Wen Jiang

## Abstract

**Background:** The Microtubule-Associated Protein Tau (MAPT) is one of the proteins that are central to neurodegenerative diseases. The nature of intracellular tau aggregates is determined by the cell types whether neuronal or glial, the participating tau isoforms, and the structure of the amyloid filament. The transmembrane protein 106B (TMEM106B) has recently emerged as another significant player in neurodegeneration and aging. In the central nervous system, the composition of the gray and white matter differs considerably. The gray matter consists of nerve cell bodies, dendrites, unmyelinated axons, synaptic terminals, astrocytes, oligodendrocytes (satellite cells) and microglia. The white matter differs from the gray for the presence of axonal tracts as the only neuronal component and for the absence of nerve cell bodies, dendrites and synaptic terminals. Cryogenic electron microscopy (cryo-EM) studies have unveiled the structure of tau and TMEM106B, from the cerebral cortex, in several neurodegenerative diseases; however, whether tau and TMEM106B filaments from the gray and white matter share a common fold requires additional investigation.

**Methods:** We isolated tau and TMEM106B from the cerebral cortex and white matter of the frontal lobes of two individuals affected by multiple system tauopathy with presenile dementia (MSTD), a disease caused by the *MAPT* intron 10 mutation +3. We used immunostaining, biochemical, genetics and cryo-EM methods to characterize tau and TMEM106B.

**Results:** We determined that tau filaments in the gray and the white matter of MSTD individuals can induce tau aggregation and have identical AGD type 2 folds. TMEM106B amyloid filaments were also found in the gray and white matter of MSTD; the filament folds were identical in the two anatomical regions.

**Conclusions:** Our findings show for the first time that in MSTD two types of amyloid filaments extracted from the gray matter have identical folds to those extracted from the white matter. Whether in this genetic disorder there is a relationship in the pathogenesis of the tau and TMEM106B filaments, remains to be determined. Furthermore, additional studies are needed for other proteins and other neurodegenerative diseases to establish whether filaments extracted from the gray and white matter would have identical folds.

## BACKGROUND

The Microtubule-Associated Protein Tau (MAPT) and the transmembrane lysosomal protein 106B (TMEM106B) belong to a group of proteins that, in the process of neurodegeneration, become misfolded and give rise to amyloid filaments whose structures are being studied at the molecular level by cryo electron microscopy (cryo-EM). The nature of intracellular tau aggregates is determined by cell type, the participating tau isoforms, and the structure of the amyloid filament. The role of TMEM106B in neurodegeneration and aging has more recently become evident following the discovery of the presence of TMEM106B amyloid filaments deriving from its carboxy terminus.

Alternative mRNA splicing of exons 2, 3, and 10 of the *MAPT* gene yields six tau isoforms in the adult human brain. These six isoforms differ due to the absence or presence of one or two acidic N-terminus inserts, and whether they contain three or four repeats (3R, 4R) of a conserved tubulin binding motif at the C-terminus [1]. The accumulation of either 3R, 4R or a mixture of 3R and 4R tau aggregates leads to multiple clinicopathological phenotypes of frontotemporal lobar degeneration associated with *MAPT* mutations. Tauopathies caused by intronic *MAPT* splice-site mutations (+3 and +16) disrupt a stem-loop structure in the mRNA, which leads to elevated levels of 4R tau isoforms in the brain [2, 3].

Multiple system tauopathy with presenile dementia (MSTD), caused by the *MAPT* intron 10 mutation +3 (IVS10+3), is characterized by superior gaze palsy, disinhibition, bradykinesia, and progressive dementia [2, 4, 5]. Neuropathologically, abundant 4R filamentous tau deposits are found in both neurons and glial cells throughout the central nervous system. Tau inclusions in neurons, astrocytes (tufted astrocytes and astrocytic plaques), and oligodendrocytes (coiled bodies) are found throughout the gray matter, while tau inclusions in astrocytes and oligodendrocytes are numerous throughout the white matter [4, 5]. The structure of tau amyloid filaments isolated from the frontal cortex of MSTD were found to be identical to the argyrophilic grain disease (AGD) type 2 fold. The AGD type 2 fold has a four-layered ordered structure accommodating amino acids 279–381 of tau, packing two protofilaments with C2 symmetry [6].

The transmembrane protein 106B (TMEM106B) is known to be associated with risk in neurodegenerative diseases and specifically frontotemporal dementias; however, the relationship between TMEM106B and tau is not well understood [7, 8]. At the cellular level, there is evidence that this protein is present in neurons, oligodendrocytes, astrocytes and microglia [8, 9]. By cryo-EM, TMEM106B filaments were recently identified in postmortem brain tissue of individuals affected by neurodegenerative diseases and of individuals with no evidence of neurodegeneration [10-13]. The TMEM106B protofilament core, spanning residues 120-254, consists of 17-19 β-strands. Amino acids 120-254 are in the C-terminus of the TMEM106B protein, situated in the lysosomal lumen [10]. The same core gives rise to filaments made of one or two protofilaments. Filaments with one protofibril (fold I) were the most common, without a noticeable relationship between fold and disease [10].

The central aims of this study were to determine whether the structure of tau and TMEM106B filaments isolated from the white matter differs from those of filaments isolated from the gray matter. In addition, we carried out neuropathologic, biochemical, and molecular genetic analyses of the two MSTD patients. Comparisons of the tau and TMEM106B filaments structures in each brain region may provide novel insights on the formation of aggregates in different cellular elements.

## METHODS

### Neuropathology

Tissue samples of gray matter and white matter were obtained from the frontal lobe. Eight-μm thick brain sections were used. For immunohistochemistry of tau, primary antibodies were AT8 (Thermo Fisher Scientific MN1020, 1:300), RD3 (Merck 05–803, 1:3,000), and RD4 (Upstate, 1:100). For immunohistochemistry of TMEM106B inclusions, we used anti-TMEM239 (residues 239–250) [10]. The signal from the antibodies was visualized using avidin–biotin followed by horseradish peroxidase-conjugated streptavidin and the chromogen diaminobenzidine. Immunohistochemical sections were counterstained with hematoxylin to show nucleus and cytoplasmic structure. In addition, the Gallyas silver method was used to demonstrate cytoplasmic inclusions, specifically of 4R tau, in nerve cells, astrocytes and oligodendroglia.

### Sanger DNA analysis

Genomic DNA was extracted from frozen brain tissue. Polymerase chain reaction (PCR) was carried out for the identification of the haplotypes of the TMEM106B gene. Oligonucleotide primer pairs 5’-GGTTTAATTTTCTTTGACATTTTGG-3’ and 5’-GGCTCAAGCAGTCCACTGAG-3’ for rs3173615 and 5’-TTGATTGTAGGGGATACAATGATG-3’and 5’-GAGTACAGGGCTTCCCAACA-3’ for rs1990622, rs1990621, and rs1990620 were used for direct sequencing, as described [10]. For rs147889591, exon 3 and flanking intronal sequences (347bp) were amplified by PCR using oligonucleotides 5’-TGTTGCTGCTGATATATGGGG-3’ and 5’-ACATCTCTTCTCAATGGGGAC-3’. The resulting amplicon was screened by agarose gel electrophoresis and directly sequenced using dye terminator sequencing.

### Whole-exome sequencing (WES)

Target enrichment made use of the SureSelectTX human all-exon library (V6, 58 megabase pairs; Agilent) and high-throughput sequencing was carried out using a HiSeq 4,000 (sx75 base-pair paired-end configuration; Illumina). Bioinformatics analyses were carried out as described [14]. In addition to the TMEM106B gene, the coding regions of genes corresponding to proteins reported to interact with the TMEM106B protein were screened for variants. These genes include *APOE, CHMP2B, CTSD, GRN, KCNMA1, KCNMB2, LAMTOR1, MAP6, MYLK, NSF, PLD3, RFC5, STX7, TARDP, TMEM106A, TMEM106B, TMEM106C, TMEM192, VAMP7*, and *VPS11* [8].

### Extraction of tau and TMEM106B filaments

Sarkosyl-insoluble fractions were prepared from the cerebral cortex and the deep white matter of freshly frozen frontal lobes as previously described [15]. Briefly, ∼4 g of gray and white matter tissues were homogenized in A68 extraction buffer consisting of 10 mM Tris–HCl, pH 7.4, 0.8 M NaCl, 1 mM EGTA, 5 mM EDTA, and 10% sucrose with protease and phosphatase inhibitors. Samples were centrifuged at 20,000 × g and the supernatants brought to 1% sarkosyl. Supernatants were incubated at room temperature (RT) while shaking. After centrifugation at 100,000 × g/1 h/4 °C, the sarkosyl-insoluble pellets were resuspended in 10 μl/g tissue 50 mM Tris–HCl, pH 7.4. The pellet was diluted in A68 extraction buffer and centrifugated at 20,000 × g/30 min/4 °C. The supernatant was centrifuged at 100,000 × g/1 h/4 °C and the final pellet resuspended in 20 mM Tris–HCl, pH 7.4 with 100 mM NaCl, and stored at 4 °C.

### Western blot analysis of tau

Samples were resolved on 10% Tris-glycine gels (BioRad) and blocked in 5% milk in TBS plus 0.1% Tween 20. Primary antibodies were diluted in TBS plus 0.1% Tween 20 at the following dilutions: AT8 (Thermo Fisher, 1:1000) and HT7 (Thermo Fisher, 1:1000).

### Immuno-electron microscopy (EM) of tau

For immuno-EM, samples were analyzed as previously described [16, 17]. AT8 antibodies were diluted 1:50 in 0.1% gelatin in PBS. Secondary antibodies used were 6 nm anti-mouse immunogold particles (Electron Microscopy Sciences). Negative staining was carried out as previously described [17]. Images were taken on a Tecnai G2 Spirit Twin scope equipped with an AMT CCD Camera.

### Tau seeding assay

Tau seeds were prepared as per filament extraction protocol. Prior to addition to cells, the resuspended pellets were sonicated with a probe sonicator at 75 W, for a total of 25 × 500 ms pulses. The probe was cleaned with isopropanol and water between samples. HEK 293 T cells stably expressing the repeat domain of Tau with a P301S mutation, fused at the C-terminus to either CFP or YFP, were obtained from ATCC [18]. Cells were cultured in DMEM supplemented with 10% FBS, 1% pen/strep, and 1% GlutaMax (Invitrogen). Cells were plated at a density of 50,000 cells/well in a 12-well plate, onto coverslips pretreated with 0.1 mg/ml poly-d-lysine for fixed cell imaging. Cells were incubated overnight at 37 °C and 5% CO_2_. The following day, 1 µl of sonicated seed material was combined with Lipofectamine 2000 (Thermo Fisher) and OptiMem medium, and incubated for 20 min at room temp. This transduction complex was then added onto the biosensor cells. Cells were incubated as before for 48 h. Cells were then washed with 1X PBS, fixed in 4% paraformaldehyde in PBS, stained with DAPI, and mounted onto microscope slides for imaging. Images were merged and cropped using ImageJ [19].

### Mass spectrometry sample preparation

Samples were diluted in 8 M urea, 50 mM Tris-HCl pH 8.5 (100 µl), reduced with 5 mM Tris (2-carboxyethyl) phosphine hydrochloride (TCEP-HCl) for 30 min at 37°C and alkylated with 10 mM chloroacetamide at RT in the dark, for 30 min. Samples were digested in two steps with LysC/trypsin (Promega). After overnight trypsin digestion in 2 M Urea, the samples were applied to Pierce detergent removal spin columns (Thermo Scientific) and then desalted on SepPak 18 cartridge (Waters Corporation) washed with 1 mL of 0.1% trifluoroacetic acid (TFA), eluted in 600 µL of 70% Acetonitrile (ACN)/0.1 % formic acid (FA)), and dried by speed vac.

### Liquid chromatography (LC) with tandem mass spectrometry (MS)

Samples were reconstituted in 50 µL of 0.1 % FA and 7 µL were injected on an Easynano LC1200 coupled with Aurora 25cm column (IonOptiks) in sonation column oven (40 °C) on an Eclipse Orbitrap mass spectrometer (Thermo Fisher Scientific). Peptides were eluted on a 115 minute gradient from 5-35 % B, increasing to 95% B over 10 minutes, and decreasing to 5% B for 5 minutes (Solvent A: 0.1% FA; Solvent B: 80% ACN, 0.1% FA). The instrument was operated with FAIMS pro 4 CVs (−30, −45, −55, −65 V), positive mode, 0.6 second cycle time per CV with APD and Easy-IC on. Full scan 400-1500 m/z with 60000 resolution, standard AGC and auto max IT, 40% RF lens, 5e4 intensity threshold, charge states 2-8, 30 sec dynamic exclusion with common settings. MS2 parameters of 1.6 m/z quadrupole isolation, 30% fixed HCD CE, 15000 orbitrap resolution, standard AGC and dynamic IT.

### MS data analysis

Raw files were loaded into PEAKS X Pro Studio 10.6 Build 20201221 (Bioinformatics Solutions) precursor ion tolerance was 10 ppm 0.02 Da, specific trypsin digestion, database searches of the reviewed Uniprot_Swissprot Homo sapiens database and common contaminants (20437 entries) with variable post-translational modifications (PTMs). PEAKS PTM and SPIDER searches were enabled to search all de novo peptides above 15% score for over 300 potential PTMs and mutations. A 0.1 % peptide FDR cutoff (−10lgP≥21.8), PTM Ascore > 10, mutation ion intensity >1 and de novo only score > 80% were applied to the data followed by PEAKS LFQ analysis. The bio-informatic analysis Gene Ontology of identified proteins was done by DAVID Bioinformatics Resources 6.8 (https://david.ncifcrf.gov). The Venn diagram was generated using BioVenn (http://www.biovenn.nl/).

### Tau and TMEM106B high-resolution cryo-EM imaging

Cryo-EM grids of gray and white matter extracts of the two MSTD patients were prepared in a biosafety level 2 cabinet. 2-3 µl of the sample were applied on a graphene oxide coated EM grid, then washed with 10 mM Tris pH 7.8 before vitrifying using a semi-automated Gatan CP3 cryo-plunger. High resolution cryo-EM movies were collected on a FEI Titan Krios at 300 kV with a Gatan K3 detector mounted on a Quantum energy filter with 20 eV slit width (Table S1).

### Helical reconstruction

CryoSPARC [20] was used to obtain motion corrected movies during data collection. The remaining steps of data processing were carried out in RELION 3.1 [21]. Filaments were manually picked and initially extracted with a box size of 1024 pixels and down-scaled to 256 pixels, to facilitate identification of filament types. Then several rounds of 2D classification were carried out to find a homogeneous subset for different types of filaments. Helical rise was estimated from the first layer line centered on the meridian in the power spectrum of the 2D classes. Helical twist was estimated from the observed cross-over distance along the filaments. Iterative 3D classification and 3D refinement using a bare cylinder as the initial reference were carried to yield the final density map. The overall resolution was calculated from gold standard Fourier shell correlations at 0.143 between two independently refined half-maps using the trueFSC.py program in JSPR [22].

### Density map and atomic model analysis

The final map was sharpened using phenix.auto_sharpen [23] and a central region (∼15 Å) of the filament was extracted for modeling using e2proc3d.py of the EMAN2 suite [24]. To distinguish polarity and handedness of the model, we manually built polyalanine models using COOT [25] in both directions, in two maps of opposite handedness resulting in four polyalanine models. To identify the protein sequence corresponding to the electron density map, phenix.sequence_from_map [26] was run with each of the four map/model pairs against all annotated sequences (8171 sequences) known to be expressed in the human brain (https://www.uniprot.org/uniprot/?query=tissue%3Abrain+reviewed%3Ayes+organism%3A“Homo+sapiens+%28Human%29+%5B9606%5D”&sort=score). In addition to these sequences, the 18 isoforms of tau (nine splicing and nine hypothetical isoforms; https://www.uniprot.org/uniprot/P10636#sequences) were also included in the analysis. The final model was refined to acceptable stereochemical parameters (Table S1) using phenix.real_space_refine [27].

### Data availability

Cryo-EM maps have been deposited in the Electron Microscopy Data Bank (EMDB) under accession numbers EMD-25995 and EMD-28943. Refined atomic models have been deposited in the Protein Data Bank (PDB) under accession numbers 7TMC and 8F9K. Mass spectrometry raw data are available at MassIVE under accession number MSV000090845. Whole-exome sequencing data have been deposited in the National Institute on Aging Alzheimer’s Disease Data Storage Site (NIAGADS; https://www.Niagads.org), under accession number NG00107.

## RESULTS

### Tau and TMEM106B in gray and white matter of individuals with MSTD

Clinical information of cases #1 and #2 have been previously reported [6]. Case #1 was a 54-year-old female and case #2 was a 63-year-old female at the time of death. Both individuals had the intronic *MAPT* +3 splice-site mutation, a G-to-A transition in the intron following exon 10. In the gray matter of the frontal cortex of both cases, tau-immunoreactive neurons and neuropil threads were seen using antibodies against tau phosphorylated at Ser202/Thr205 (AT8) (Fig. 1a,c) and 4R tau (RD4) (SFig. 1). In the subcortical white matter, numerous AT8 and RD4 immunoreactive intracytoplasmic inclusions in oligodendroglia (coiled bodies) were present. Coiled bodies were strongly positive for Gallyas silver staining (SFig. 1). Intracellular 3R immunoreactive deposits were not present in either gray or white matter (SFig. 1).

**Figure 1.**
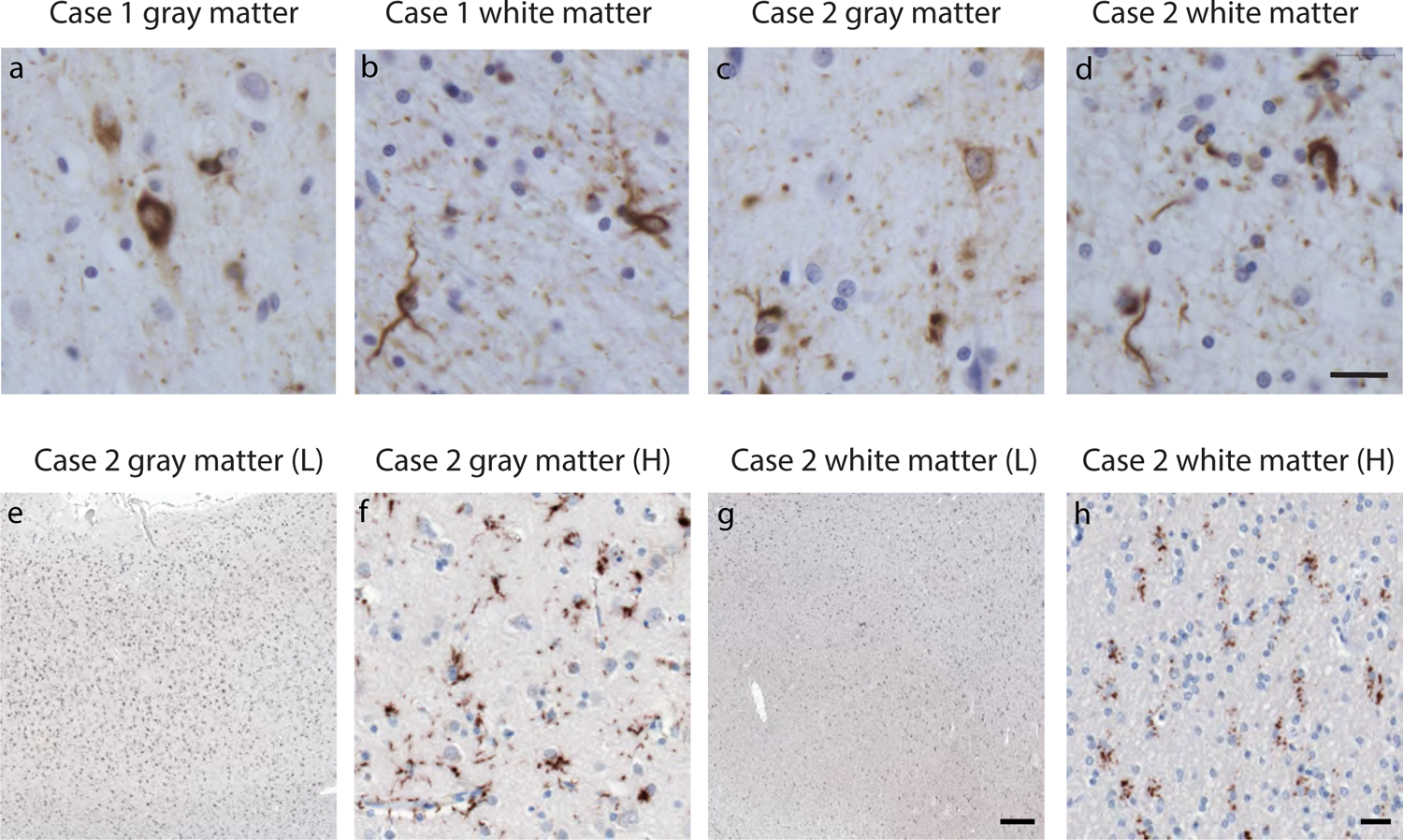
Photomicrographs showing neuronal and glial inclusions labeled with Tau and TMEM106B antibodies. Nerve cell bodies, glial cells, and dot-like neuropil elements labeled for phosphorylated tau in the gray matter (a,c). Oligodendroglial cells and their processes with intracytoplasmic inclusions, some appearing as coiled bodies, labeled for phosphorylated tau in the white matter (b,d). At low magnification, cellular profiles containing inclusions labeled for TMEM106B antibodies in the gray and white matter (e,g). At high magnification, inclusions labeled for TMEM106B in the cytoplasm of neuronal and glial cell body as well as glial cytoplasmic processes in the gray matter (f); inclusions labeled for TMEM106B in oligodendroglial of the white matter (h). Anti-tau antibody AT8 (a,b,c,d). Anti-TMEM106B antibody TMEM239 (239-250) (e,f,g,h). Scale bar: 20 µm (a,b,c,d,f,h). Scale bar: 200 µm (e,g).

Immunohistochemistry of gray and white matter using anti-TMEM239 (residues 239– 250) showed numerous cellular profiles containing immunopositive inclusions in the cytoplasm of the cell body and in elongated cytoplasmic processes in both cerebral cortex and subcortical white matter (Fig. 1e-h). In the gray matter, the most numerous immunopositive cells appear to be astrocytes. Neurons appeared to have fewer immunopositive inclusions; however, considering the severe neuronal atrophy and loss that had occurred in the frontal cortex, it was difficult to clearly recognize the identity of some cells. Their identification was possible based on the morphology of the nucleus and the presence of immunopositive inclusions that were occasionally globular in shape and extended from the cytoplasm of the cell body to multiple cellular processes. Some immunopositive processes were seen near the vascular endothelium suggesting their astrocytic nature. In some instances, the immunopositive profiles appeared as small, dotted inclusions. In the white matter, immunopositive glial cells were less numerous than those observed in the gray matter. The lack of neurons in the white matter facilitated the recognition of the cellular identity. Both oligodendroglial cells and astrocytes appeared to be immunopositive with the TMEM239 antibody.

### Biochemistry of tau from gray and white matter samples

Western blot analysis of the sarkosyl-insoluble fractions using antibodies to human tau (HT7) and phosphorylated tau (AT8) showed the presence of tau bands with a migration pattern corresponding to 4R tau, with identical electrophoretic mobility on either gray or white matter of both cases (Fig. 2a). Immunogold transmission electron microscopy preparations of dispersed tau filaments from gray and white matter showed twisted ribbon-like filaments that were decorated by the AT8 antibody, supporting the concept that the filaments were phosphorylated (Fig. 2b). Mass spectrometric analysis of the sarkosyl-insoluble fractions extracted from the gray and white matter of cases #1 and #2 revealed the presence of numerous post-translational modifications (PTMs) on tau, with no significant difference between gray and white matter or between cases. A number of proteins that co-purified with tau were identified by mass spectrometry and constitute the MSTD-tau “interactome” (SFig. 1m). A total of 2,006 proteins were found in association with tau deposits in the gray and white matter of both cases. Case #1 had 378 unique proteins in the gray matter, 339 proteins in the white matter, and 131 common proteins in the gray and white matter. Case #2 had 353 unique proteins in the gray matter, 377 proteins in the white matter, and 509 common proteins in the gray and white matter. The purified sarkosyl-insoluble fractions enriched for tau aggregates induced tau aggregation in a biosensor cell system. The tau isolated from the gray matter led to the formation of aggregates similar to those from tau isolated from the white matter of both cases. Tau isolated from the frontal cortex of a normal control or treatment with lipofectamine alone did not lead to the formation of aggregates (Fig. 2c).

**Figure 2.**
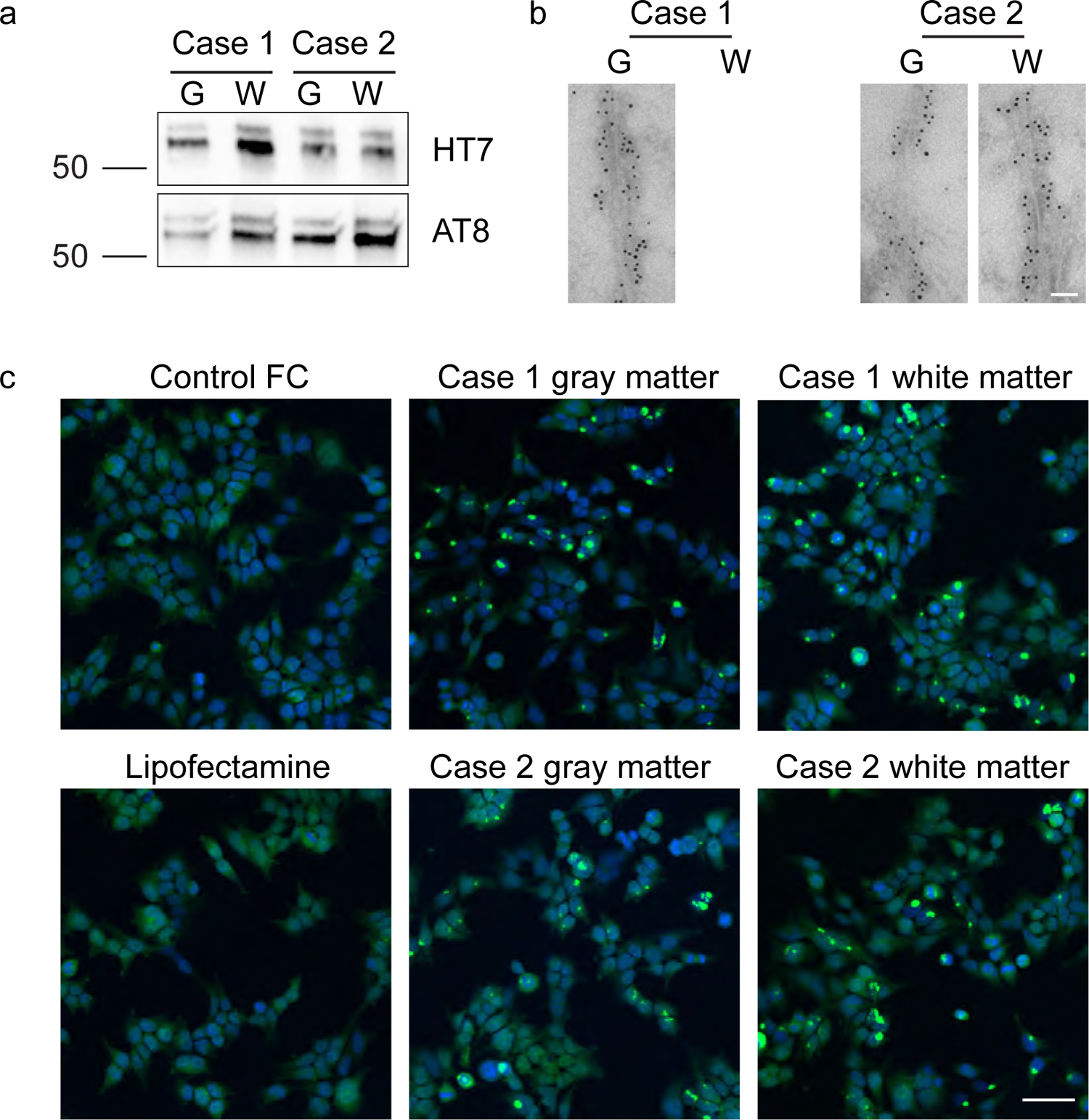
Immunoblotting, negative-stain immunoelectron microscopy and tau biosensor imaging of tau filaments from gray and white matter. a) Western blots of sarkosyl-insoluble tau fractions from MSTD (Cases #1 and #2) show that tau aggregates consist of hyperphosphorylated tau with an identical electrophoretic pattern, consistent with 4R-tau, in samples from gray (G) and white (W) matter using phosphorylation independent (HT7) and dependent (AT8) antibodies. b) Representative images of immuno-EM of sarkosyl insoluble tau filaments labelled by AT8. c) Tau biosensor cells incubated with the sarkosyl-insoluble fraction obtained from gray and white matter from MSTD (Cases #1 and #2). Control brain frontal cortex (FC) and lipofectamine only (no brain) were also analyzed. Gray and white matter from cases #1 and #2 show tau seeding activity, whereas the control brain and lipofectamine alone do not seed. Scale bars: b) 50 nm; c) 50 µm.

### Cryo-EM of tau filaments from gray and white matter

We determined the structure of tau filaments at high resolution by cryo-EM imaging and 3D reconstruction. Visual inspection of cryo-EM micrographs showed the presence of different types of filaments in gray and white matter of both cases (SFig. 2a). 2D classification of filaments in RELION revealed that all filaments had a cross over distance of ∼2,000 Å (SFig. 2b). We identified a group of thinner filaments that had a width of 130-150 Å and a group of thicker filaments that had a larger variation in width (130-350 Å). The power spectrum of these 2D classes (SFig. 2c) displayed a layer line at ∼4.8 Å with a peak on the meridian, suggesting that the filaments have around ∼4.8 Å helical rise/subunit without 2_1_ screw symmetry. Helical reconstruction in RELION revealed three distinct kinds of cross-β filaments in each of the four samples (SFig. 2b). Using RELION 3D classification, tau filaments were resolved using the following helical parameters, twist 0.41° and rise 4.8 Å. Filaments were made up of a doublet form of the filaments and were further refined to a resolution of ∼4.5 Å (Fig. 3a). The doublets were composed of two protofilaments with C2 symmetry corresponding to the AGD type 2 filaments [6]. The AGD type 2 filaments have a four-layered ordered fold comprising residues 279–381 of tau (Fig. 3b). Type 1 and 3 AGD filaments were not observed. Filaments from gray matter were identical to filaments from white matter in both MSTD patients.

**Figure 3.**
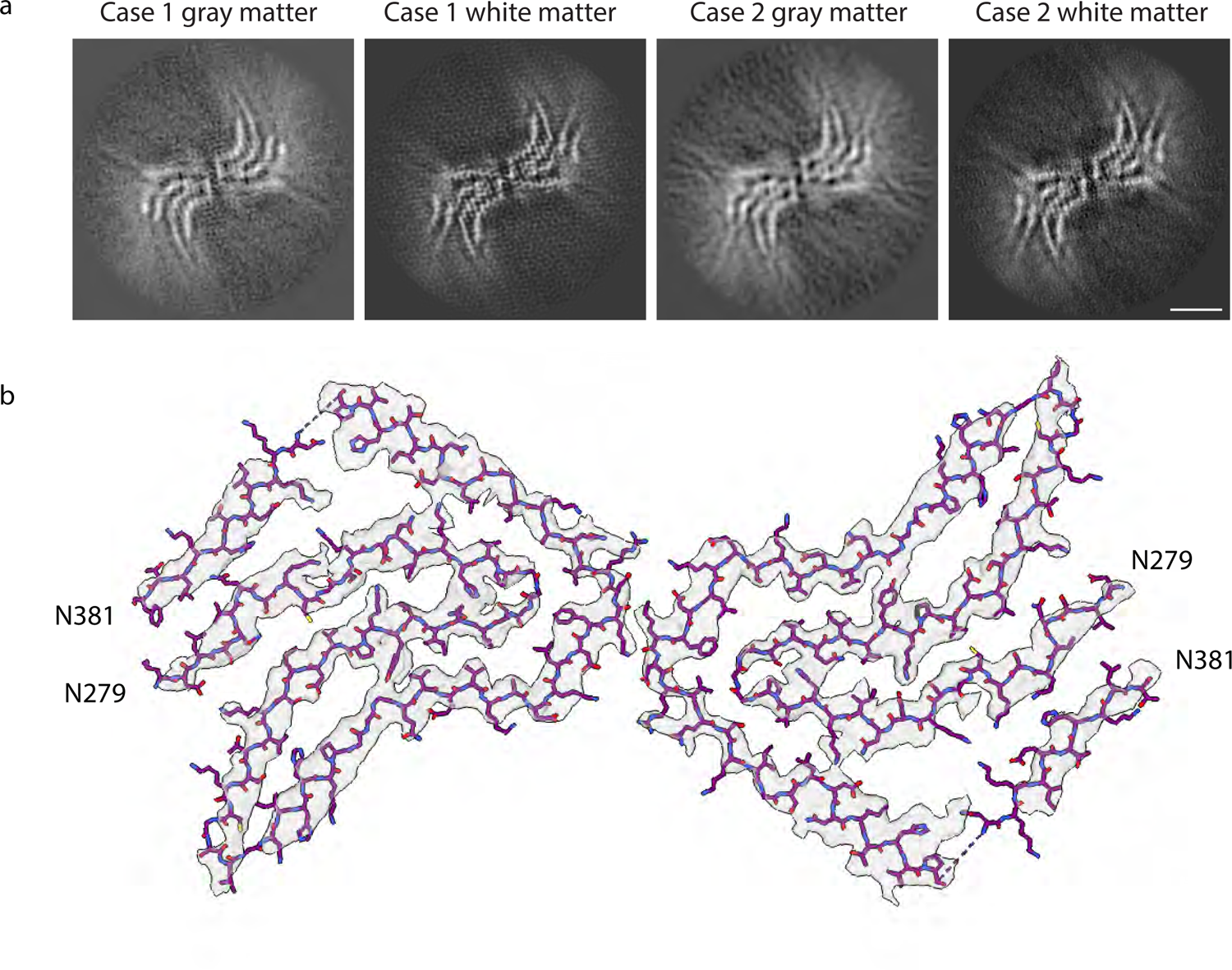
Cryo-EM structure of tau filaments. a) Cryo-EM maps for tau filaments from gray and white matter from MSTD cases #1 and #2 showing the AGD type 2 filaments. b) Cryo-EM density map and atomic model of AGD type 2 filaments. Scale bar 5 nm.

### Cryo-EM of TMEM106B filaments from gray and white matter

Helical reconstruction in RELION of the cross-β filaments (SFig. 2) revealed two additional types of filaments, a singlet and a doublet form of the same fold with helical parameters twist of 0.41° and rise of 4.8 Å (Fig. 4a). The singlet and doublet forms were resolved to 2.9 and 3.5 Å resolution, respectively. The doublet form has C2 symmetry, and it comprises two identical protofilaments of the singlet type. *Ab initio* model building efforts with phenix.map_to_model, buccaneer, deepTracer or manual model building in COOT revealed that the density map could not be explained with any of the isoforms of tau. Manually built polyalanine models in two different directions were used to identify and score the density at each Cα position using phenix.sequence_from_map and to unambiguously assign the TMEM106B sequence to the electron density (SFig. 3). We observed extra densities which corresponded to glycosylation at residues N145, N151, N164 and N183 of TMEM106B (Fig. 4). The singlet and doublet filaments of TMEM106B displayed the same fold, consisting of 17 β strands formed by residues Ser120 to Gly254 (Fig. 4), as previously described [10]. Each chain is stacked along the helical axis and these successive layers interact through main chain hydrogen bonds reminiscent of parallel β-strand packing. Within a chain, strands β1, β14, β15, and β16 make up the central core. The packing of the β-strands is further strengthened by the presence of a disulfide bond between Cys214 and Cys253. Interestingly, the N-terminal residue Ser120 was hidden in the interior of the fibril and away from the exposed solvent (Fig. 4b, SFig. 4). The remaining residues of the C-terminus Arg255 to Gln274 were likely to be disordered and hence not resolved in the electron density. In the doublet filaments of TMEM106B, there appears to be no direct interaction between the two protofilaments. Instead, the protofilament interactions appear to be mediated through a non-proteinaceous density at the two-fold axis (Fig. 4). This density was surrounded by a pair of arginine and lysine residues (Lys178 and Arg180) from one chain and its dyad axis related monomer suggesting a negatively charged factor mediating the interprotofilament packing (Fig. 4, SFig. 4, 5). Mass spectrometric analysis of the sarkosyl-insoluble fraction extracted from the gray and white matter of case #2 determined the presence of three tryptic peptides corresponding to residues 130-SAYVSYDVQK-139, 130-SAYVSYDVQKR-140, and 248-YQYVDCGR-255. Residue Arg255 is not included in the fibril core and may be part of the C-terminal fuzzy coat of TMEM106B.

**Figure 4.**
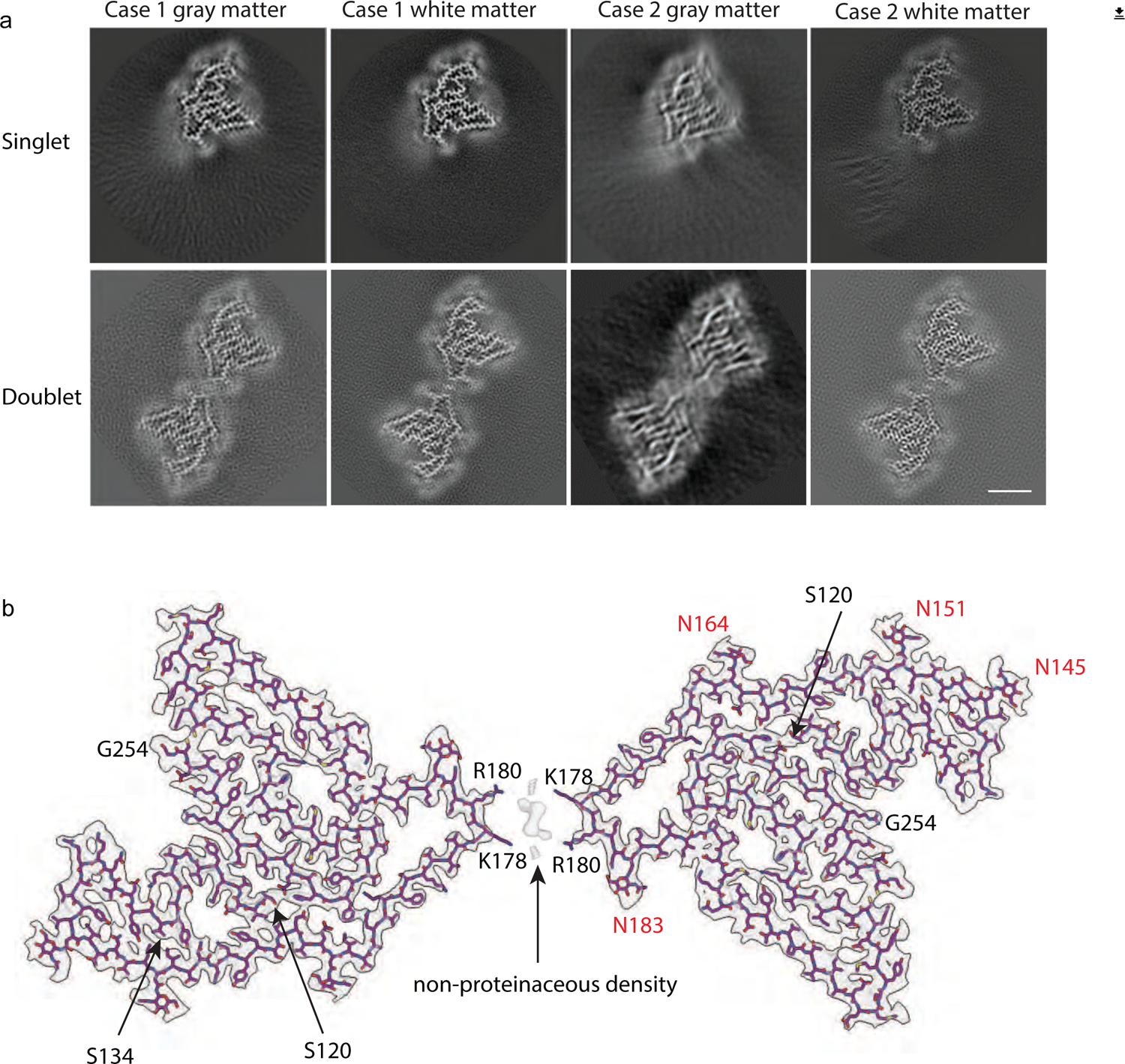
Cryo-EM structure of TMEM106B filaments. a) Cryo-EM maps for TMEM106B filaments from gray and white matter from MSTD cases #1 and #2 showing filaments made of a single protofilament and filaments comprising two protofilaments. Scale bar: 5 nm. b) Cryo-EM density map and atomic model of the doublet form of a TMEM106B filament. The N-terminus (S120) and C-terminus (G254) are indicated in black. Asn (N) glycosylated residues are indicated in red. K178 and R180 are involved in the binding of an unknown cofactor. Position of Ser134 (S134) that is replaced in one allele by Asn (S134N) in Case #1 is indicated.

### TMEM106B and interacting partners’ genetic analyses

Genotyping of TMEM106B in both cases showed that MSTD individuals were homozygous for Thr185. Homozygosity at codon 185 was confirmed by Sanger sequencing. WES analysis revealed a point mutation, c.401g>a, resulting in an asparagine replacing serine at residue 134 (S134N) in one allele of the *TMEM106B* gene in Case #1 (Table S2). The change (rs 147889591) was confirmed by direct sequencing. We also analyzed a number of TMEM106B interacting partners [8]. A total of 20 genes were analyzed. Additional variants were found in *APOE, TMEM106C, VPS11, PLD3, and KCNMA1*. Coding variants were also present in *APOE, TMEM106C, VPS11*, and *KCNMA1* in Case #2 (Table S2).

## DISCUSSION

In recent years, extensive cryo-EM studies have unveiled the structure of amyloid filaments involved in neurodegeneration. This work led to the discovery of the complexity of the structure of filaments formed by proteins such as tau, α-synuclein, amyloid-β, prion protein amyloid, TDP-43, and more recently, TMEM106B. It has been shown that filaments of these proteins extracted from the gray matter differ according to the specific disease process in which they are involved [6, 17, 28-34].

TMEM106B is a lysosomal transmembrane protein vital for lysosomal health and has been implicated as a risk factor and/or disease modulator in many neurodegenerative diseases [8]. Recently, while investigating *ex-vivo* filaments from individuals affected by neurodegenerative diseases, it was serendipitously discovered that TMEM106B filaments co-exist with other aggregated protein filaments in sporadic and inherited tauopathies, amyloid-β amyloidoses, synucleinopathies and TDP-43 proteinopathies [10-13]. Insoluble TMEM106B accumulation has recently also been reported to be associated with multiple sclerosis and has been emphasized for its role in oligodendroglia [35]. It is important to note that TMEM106B filaments have been found in the brain of elderly individuals being particularly severe in individuals 75 years and older, but not present in the brain of individuals younger than 46 years of age [10, 13]. Furthermore, TMEM106B folds seem to be similar across the different diseases unlike tau filaments. These data strongly suggest that TMEM106B filament formation is an aging-related process [10]. The mechanism of TMEM106B filament formation is unknown. Filaments may be formed within the lysosome and released or the C-terminal domain may aggregate in the cytosol once it is processed at Arg119/Ser120 and released.

It is unknown whether tau filaments in gray matter and in white matter have identical structures in the same disease even though it has been proposed that oligodendrocytes and astrocytes play an important role in tau propagation [36, 37]. A similar question can be asked about the structure of TMEM106B in the same disease. Therefore, the central aim of this study was to determine the near atomic structure of tau and TMEM106B in the white matter of individuals with MSTD and compare with the filament structure of both tau and TMEM106B isolated from gray matter.

Depending on the isoforms involved in the specific disorder, tau aggregates are found in neurons or in neurons and astrocytes in the gray matter; whereas, tau aggregates are found in oligodendrocytes and astrocytes in the white matter. We isolated sarkosyl-insoluble material from gray and white matter from two individuals affected by MSTD and determined the structures of tau to be identical within each brain region. We observed the same tau structure, corresponding to the AGD type 2 fold in both the cerebral cortex and subcortical white matter of the frontal lobe of the two individuals. The AGD type 2 fold was originally described in the gray matter from the MSTD cases #1 and #2, and the gray matter from two cases of argyrophilic grain disease (AGD), one of aging-related tau astrogliopathy (ARTAG), and three of frontotemporal dementia (FTD) with an intronic *MAPT* mutation (IVS10+16) [6]. Our data from gray versus white matter in MSTD suggest a common mechanism of tau folding that is indistinguishable between brain regions. Moreover, the sarkosyl-insoluble fractions derived from gray and white matter in MSTD induced similar seeded aggregation on a FRET-based biosensor cell line. Since the *MAPT* IVS10+3 mutation leads to overproduction of 4R tau, it remains to be determined the exact role that tau propagation plays in tau folding in different brain areas and in the potential involvement of neurons and glia in the sequence of events leading to tau misfolding and propagation *in vivo*.

TMEM106B filaments were obtained from the same isolated sarkosyl-insoluble material prepared to extract tau filaments [16]. Other studies reported TMEM106B filaments isolated using variations of the sarkosyl extraction protocol [10-13]. It remains to be seen whether small variations in the extraction protocol could facilitate or hinder the extraction of TMEM106B. We observed the presence of TMEM106B filaments with an ordered core comprising residues S120 to G254. We identified singlet and doublet filaments of TMEM106B that display the same fold, containing a five-layered core corresponding to fold I of Schweighauser et al. [10]. TMEM106B fold I has been seen to be able to accommodate threonine at codon 185 (T185), with both MSTD individuals being homozygous for T185. As previously described, we observed the presence of densities corresponding to four glycosylation sites (N145, N151, N164, and N183). Case #1 was heterozygous for a polymorphism at codon 134 (S134N) that is inside the core; however, due to the similarities in the electrodensity of both residues, it was not possible to establish by cryo-EM whether filaments contained Ser or Asn at 134. Asn134 may not be glycosylated due to the lack of a consensus sequence for N-glycosylation at 134 (Asn-Xaa-Ser/Thr (where Xaa is not Pro) in the TMEM106B sequence. By mass spectrometry analysis, we identified TMEM106B peptides in gray and white matter in MSTD. Interestingly, the tryptic peptides were identical to the peptides previously reported in FTLD-TDP [11]. The trypsin proteolysis/mass spectrometry approach provides additional insights into the structure of TMEM106B within the fibrillar polymer. Of the eleven potential trypsin-cleavage sites in the core (residues S120 to G254), only five were accessible to trypsin. Lack of cleavage at Lys124 in the β1 strand supports the atomic model generated by cryo-EM, in which the N-terminal residue Ser120 is hidden in the interior of the fibril and away from the exposed solvent. Interestingly, the Lys178 and Arg180 residues that are involved in the interactions between two protofilaments of fold I and a non-proteinaceous density at the two-fold axis were also fully protected.

This study has identified for the first time that tau filaments in gray and white matter have identical structures in individuals with MSTD, corresponding to the AGD type 2 fold. Although additional cases and diseases may need to be analyzed, our data support the notion that there is not region/cell type-specific folding of tau, and that the same tau fold may be found in neurons and glia in the same disease. Our finding of TMEM106B in gray and white matter regions of the brain imply a similar mechanism of TMEM106B fibrillization in both neuronal and glial cell types. However, considering that the frontal cortex of the two MSTD patients had undergone severe atrophy and neuronal loss, we need to be cautious in assigning the presence of the TMEM106B extracted from the frontal cortex to the two cell populations. Further research may clarify whether TMEM106B filaments can seed native TMEM106B and propagate in a prion-like manner such as tau and α-synuclein, perhaps connecting protein misfolding with the endolysosomal system in aging and neurodegeneration [38].

## Acknowledgments

We thank the family of the patients for donating brain tissue; Drs. M. Goedert and S. H. W. Scheres for anti-TMEM239 antibody. G. Qi, M. Jacobsen, and R. M. Richardson for technical support. This work was supported by the US National Institutes of Health (P30-AG010133, U01-NS110437, RF1-AG071177, R01-AG080001), and the Department of Pathology and Laboratory Medicine, Indiana University School of Medicine (to B.G. and R.V.). G.I.H was supported by K99-AG078500. K.A.O. was supported by T32-GM132024. We thank the Purdue Rosen Center for Advanced Computing for providing computing resources. We thank S. Wilson for maintaining the Jiang lab computational resources. We thank the Purdue Cryo-EM Facility (http://cryoem.bio.purdue.edu) for the use of the Titan Krios microscope. Mass spectrometry was provided by the Indiana University School of Medicine Center for Proteome Analysis. We acknowledge the Center for Medical Genomics of Indiana University School of Medicine for next-generation DNA sequencing.

## Abbreviations

AGD: argyrophilic grain disease
ARTAG: aging-related tau astrogliopathy
CNS: central nervous system
cryo-EM: cryogenic electron microscopy
FA: formic acid
FTD: frontotemporal dementia
EM: electron microscopy
EMDB: Electron Microscopy Data Bank
LC: liquid chromatography
MAPT: microtubule-associated protein tau
MS: mass spectrometry
MSTD: multiple system tauopathy with presenile dementia
PCR: polymerase chain reaction
PDB: Protein Data Bank
PTMs: post-translational modifications
TCEP-HCl: tris (2-carboxyethyl) phosphine hydrochloride
TDP-43: TAR DNA-binding protein-43
TMEM106B: transmembrane protein 106B
WES: whole-exome sequencing

## References

1. Goedert M, Jakes R. Expression of separate isoforms of human tau protein: correlation with the tau pattern in brain and effects on tubulin polymerization. EMBO J. 1990;9:4225–30.

2. Spillantini MG, Murrell JR, Goedert M, Farlow MR, Klug A, Ghetti B. Mutation in the tau gene in familial multiple system tauopathy with presenile dementia. Proc. Natl Acad. Sci. USA. 1998;95:7737–41.

3. Hutton M, Lendon CL, Rizzu P, Baker M, Froelich S, Houlden H, et al. Association of missense and 5’-splice-site mutations in tau with the inherited dementia FTDP-17. Nature. 1998;393:702–5.

4. Spillantini MG, Goedert M, Crowther RA, Murrell JR, Farlow MR, Ghetti B. Familial multiple system tauopathy with presenile dementia: a disease with abundant neuronal and glial tau filaments. Proc. Natl Acad. Sci. USA. 1997;94:4113–8.

5. Spina S, Farlow MR, Unverzagt FW, Kareken DA, Murrell JR, Fraser G, et al. The tauopathy associated with mutation +3 in intron 10 of Tau: characterization of the MSTD family. Brain. 2008;131(Pt 1):72–89.

6. Shi Y, Zhang W, Yang Y, Murzin AG, Falcon B, Kotecha A, et al. Structure-based classification of tauopathies. Nature. 2021; 598(7880):359–63.

7. Van Deerlin VM, Sleiman PM, Martinez-Lage M, Chen-Plotkin A, Wang LS, Graff-Radford NR, et al. Common variants at 7p21 are associated with frontotemporal lobar degeneration with TDP-43 inclusions. Nat Genet. 2010;42(3):234–9.

8. Perneel J, Rademakers R. Identification of TMEM106B amyloid fibrils provides an updated view of TMEM106B biology in health and disease. Acta Neuropathol. 2022;144(5):807–19.

9. Perneel J, Neumann M, Heeman B, Cheung S, Van den Broeck M, Wynants S, et al. Accumulation of TMEM106B C-terminal fragments in neurodegenerative disease and aging. Acta Neuropathol. 2022;doi:10.1007/s00401-022-02531-3.

10. Schweighauser M, Arseni D, Bacioglu M, Huang M, Lövestam S, Shi Y, et al. Age-dependent formation of TMEM106B amyloid filaments in human brains. Nature. 2022;605:310–4.

11. Chang A, Xiang X, Wang J, Lee C, Arakhamia T, Simjanoska M, et al. Homotypic fbrillization of TMEM106B across diverse neurodegenerative diseases. Cell 2022;185:1346–55.

12. Jiang YX, Cao Q, Sawaya MR, Abskharon R, Ge P, DeTure M, et al. Amyloid fbrils in FTLD-TDP are composed of TMEM106B and not TDP-43. Nature. 2022;605:304–9.

13. Fan Y, Zhao Q, Xia W, Tao Y, Yu W, Chen M, et al. Generic amyloid fbrillation of TMEM106B in patient with Parkinson’s disease dementia and normal elders. Cell Res 2022;32:585–8.

14. Farlow JL, Robak LA, Hetrick K, Bowling K, Boerwinkle E, Coban-Akademir ZH, et al. Whole-exome sequencing in familial Parkinson disease. JAMA Neurol. 2016;73:68–75.

15. Goedert M, Spillantini MG, Jakes R, Rutherford D, Crowther RA. Multiple isoforms of human microtubule-associated protein tau: sequences and localization in neurofibrillary tangles of Alzheimer’s disease. Neuron. 1989;3(4):519–26.

16. Goedert M, Spillantini MG, Cairns NJ, Crowther RA. Tau proteins of Alzheimer paired helical filaments: abnormal phosphorylation of all six brain isoforms. Neuron. 1992;8:159–68.

17. Hallinan GI, Hoq MR, Ghosh M, Vago FS, Fernandez A, Garringer HJ, et al. Structure of Tau filaments in Prion protein amyloidoses. Acta Neuropathol. 2021;142(2):227–41.

18. Holmes BB, Furman JL, Mahan TE, Yamasaki TR, Mirbaha H, Eades WC, et al. Proteopathic tau seeding predicts tauopathy in vivo. Proc Natl Acad Sci USA 2014;111(41):E4376–85.

19. Schneider CA, Rasband WS, Eliceiri KW. NIH Image to ImageJ: 25 years of image analysis. Nat Methods. 2012;9(7):671–5. doi:

20. Punjani A, Rubinstein JL, Fleet DJ, Brubaker MA. cryoSPARC: algorithms for rapid unsupervised cryo-EM structure determination. Nat Methods. 2017;14(3):290–6.

21. Scheres SHW. A Bayesian view on Cryo-EM structure determination. J Mol Biol. 2012;415(2):406–18.

22. Sun C, Gonzalez B, Vago FS, Jiang W. High resolution single particle Cryo-EM refinement using JSPR. Prog Biophys Mol Biol. 2021;160:37–42.

23. Terwilliger TC, Adams PD, Afonine PV, Sobolev OV. A fully automatic method yielding initial models from highresolution cryo-electron microscopy maps. Nat Methods. 2018;15(11):905–8.

24. Tang G, Peng L, Baldwin PR, Mann DS, Jiang W, Rees I, et al. EMAN2: an extensible image processing suite for electron microscopy. J Struct Biol. 2007;157(1):38–46.

25. Emsley P, Lohkamp B, Scott WG, Cowtan K. Features and development of Coot. Acta Crystallogr D Biol Crystallogr. 2010;66(Pt 4):486–501.

26. Terwilliger TC, Sobolev OV, Afonine PV, Adams PD, Ho CM, Li X, et al. Protein identification from electron cryomicroscopy maps by automated model building and side-chain matching. Acta Crystallogr D Struct Biol. 2021;77(Pt 4):457–62.

27. Afonine PV, Klaholz BP, Moriarty NW, Poon BK, Sobolev OV, Terwilliger TC, et al. New tools for the analysis and validation of cryo-EM maps and atomic models. Acta Crystallogr D Struct Biol. 2018;74(Pt 9):814–40.

28. Fitzpatrick AWP, Falcon B, He S, Murzin AG, Murshudov G, Garringer HJ, et al. Cryo-EM structures of tau filaments from Alzheimer’s disease. Nature. 2017; 547(7662):185–90.

29. Falcon B, Zhang W, Murzin AG, Murshudov G, Garringer HJ, Vidal R, et al. Structures of filaments from Pick’s disease reveal a novel tau protein fold. Nature. 2018;561(7721):137–40.

30. Falcon B, Zivanov J, Zhang W, Murzin AG, Garringer HJ, Vidal R, et al. Novel tau filament fold in chronic traumatic encephalopathy encloses hydrophobic molecules. Nature. 2019;568(7752):420–3.

31. Arakhamia T, Lee CE, Carlomagno Y, Duong DM, Kundinger SR, Wang K, et al. Posttranslational modifications mediate the structural diversity of tauopathy strains. Cell 2020;180(4):633–44.

32. Zhang W, Tarutani A, Newell KL, Murzin AG, Matsubara T, Falcon B, et al. Novel tau filament fold in corticobasal degeneration. Nature. 2020;580(7802):283–7.

33. Arseni D, Hasegawa M, Murzin AG, Kametani F, Arai M, Yoshida M, Ryskeldi-Falcon B. Structure of pathological TDP-43 filaments from ALS with FTLD. Nature. 2022;601(7891):139–43.

34. Hallinan GI, Ozcan KA, Hoq MR, Cracco L, Vago FS, Bharath SR, et al. Cryo-EM structures of prion protein filaments from Gerstmann-Sträussler-Scheinker disease. Acta Neuropathol. 2022;144(3):509–20.

35. Shafit-Zagardo B, Sidoli S, Goldman, JE, DuBois JC, Corboy JR, Strittmatter SM, et al. TMEM106B is increased in Multiple Sclerosis plaques, and deletion causes accumulation of lipid after demyelination. bioRxiv 2022.05.05.490697; doi: https://doi.org/10.1101/2022.05.05.490697

36. Ferrer I. Oligodendrogliopathy in neurodegenerative diseases with abnormal protein aggregates: The forgotten partner. Prog Neurobiol. 2018;169:24–54.

37. Amro Z, Yool AJ, Collins-Praino LE. The potential role of glial cells in driving the prion-like transcellular propagation of tau in tauopathies. Brain Behav Immun Health. 2021;14:100242.

38. Shao W, Todd TW, Wu Y, Jones CY, Tong J, Jansen-West K, et al. Two FTD-ALS genes converge on the endosomal pathway to induce TDP-43 pathology and degeneration. Science. 2022;378(6615):94–9.

