## Supplementary materials for "Cross-β helical filaments of Tau and TMEM106B in Gray and White Matter of Multiple System Tauopathy with presenile Dementia"

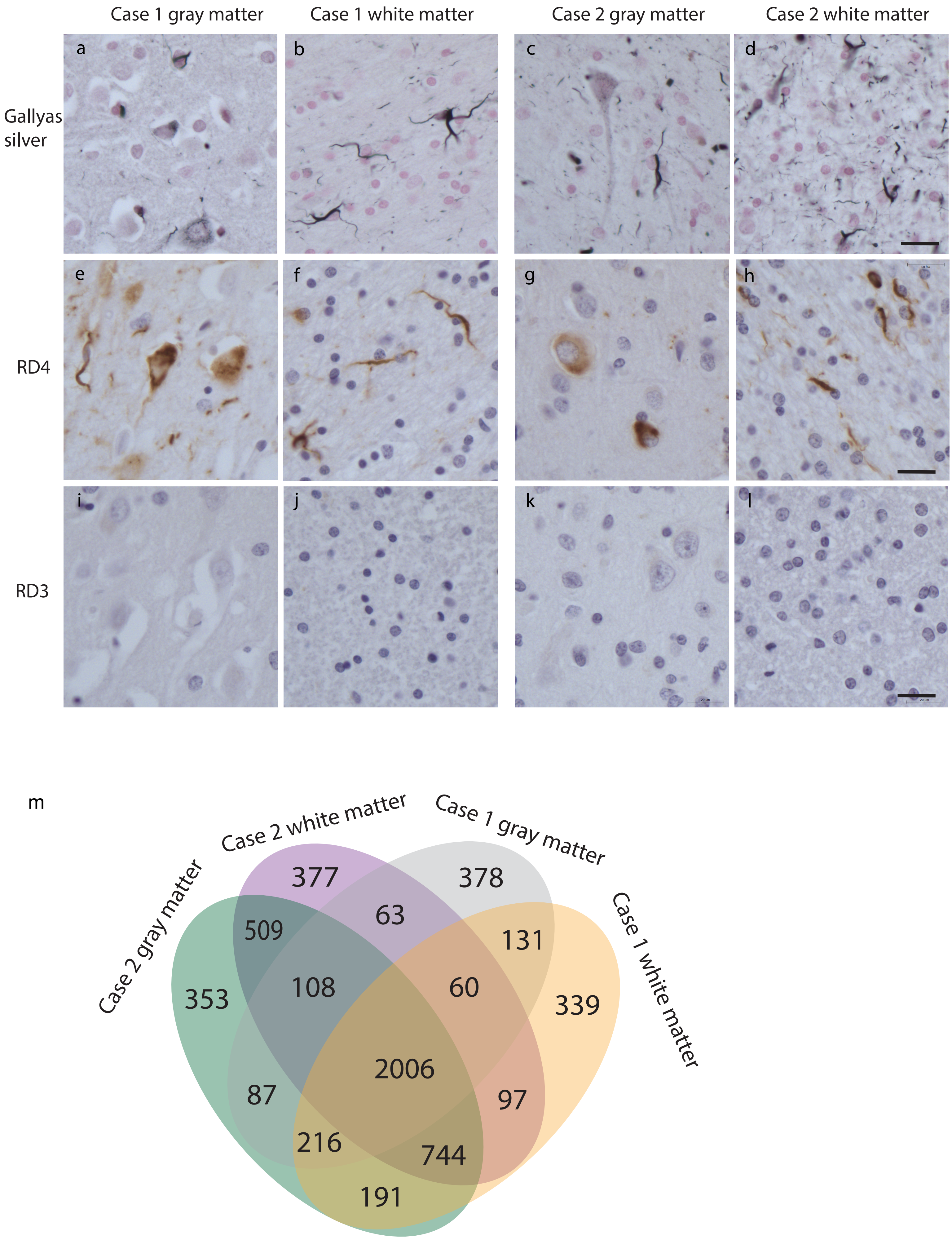
Case 1 gray matter
Case 1 white matter
Case 2 gray matter
Case 2 white matter
Gallyas silver
RD4
RD3
a
b
c
d
e
f
g
h
i
j
k
l
m
Case 2 gray matter
Case 2 white matter
Case 1 gray matter
Case 1 white matter
377
378
509
63
131
353
108
60
339
2006
87
97
216
744
191

SFigure 1

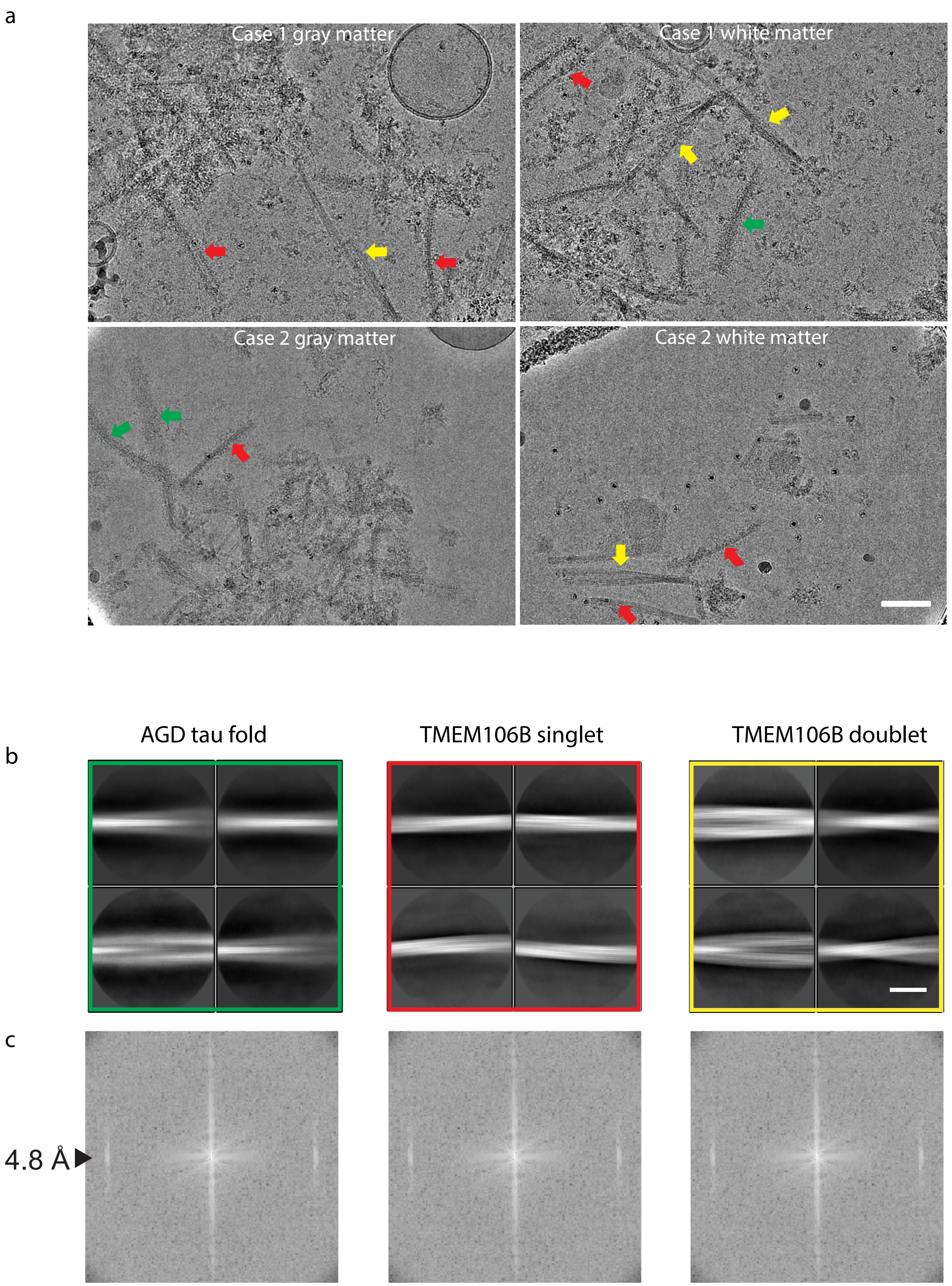
a
Case 1 gray matter
Case 1 white matter
Case 2 gray matter
Case 2 white matter
b
AGD tau fold
TMEM106B singlet
TMEM106B doublet
c
4.8 Å

a

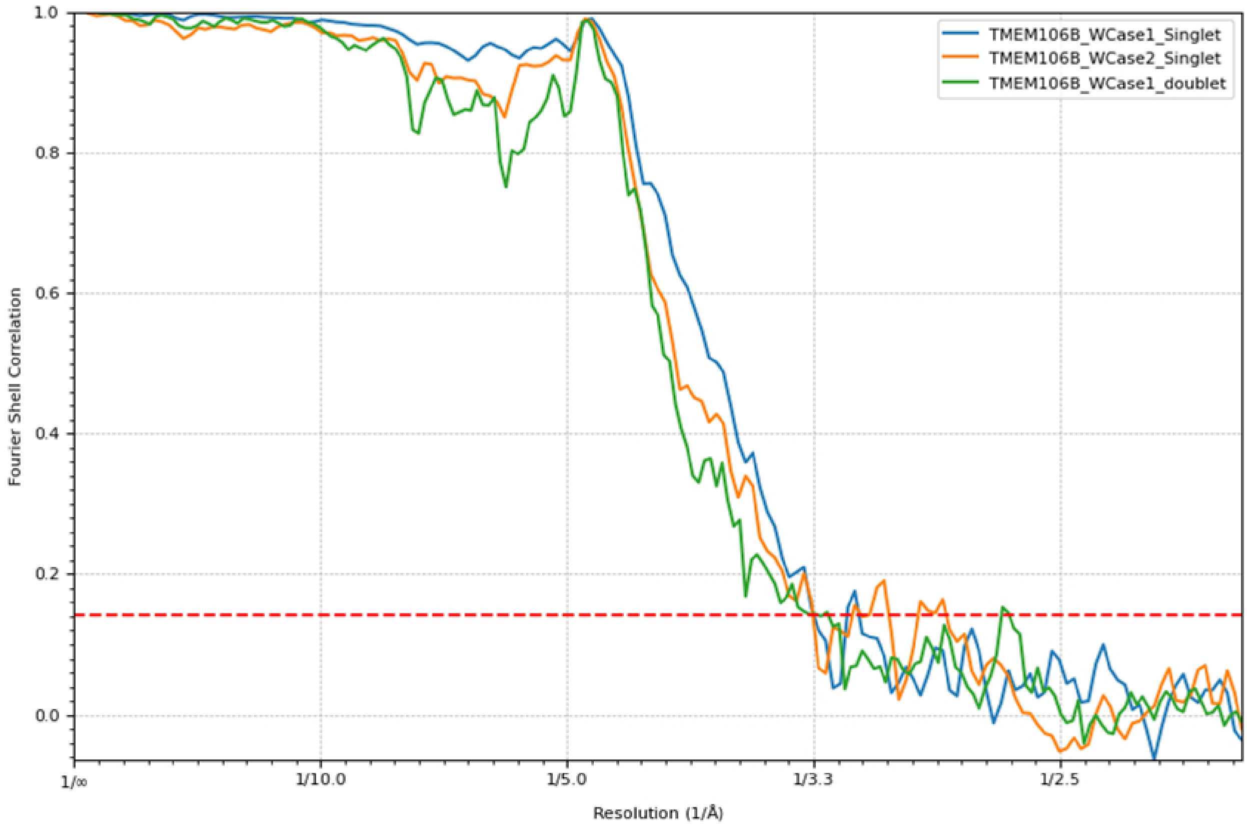
TMEM106B_WCase1_Singlet
TMEM106B_WCase2_Singlet
TMEM106B_WCase1_doublet
Fourier Shell Correlation
Resolution (1/Å)
1.0
0.8
0.6
0.4
0.2
0.0
1/∞
1/10.0
1/5.0
1/3.3
1/2.5

b

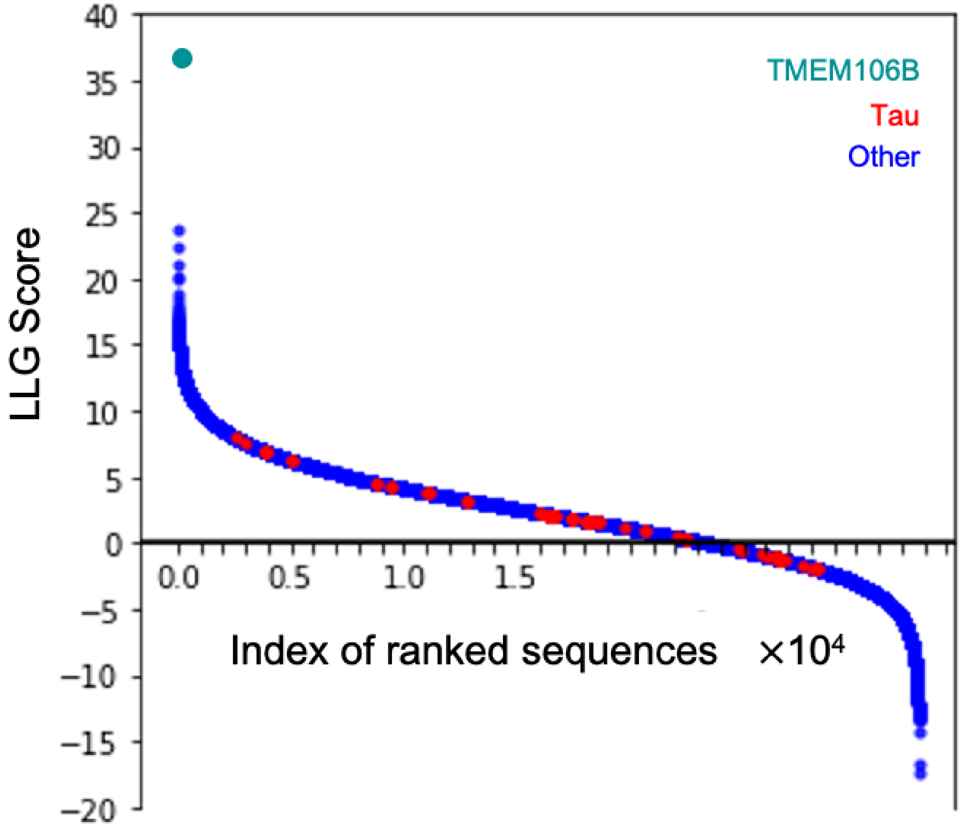
TMEM106B
Tau
Other
LLG Score
Index of ranked sequences ×10⁴
40
35
30
25
20
15
10
5
0
−5
−10
−15
−20
0.0
0.5
1.0
1.5

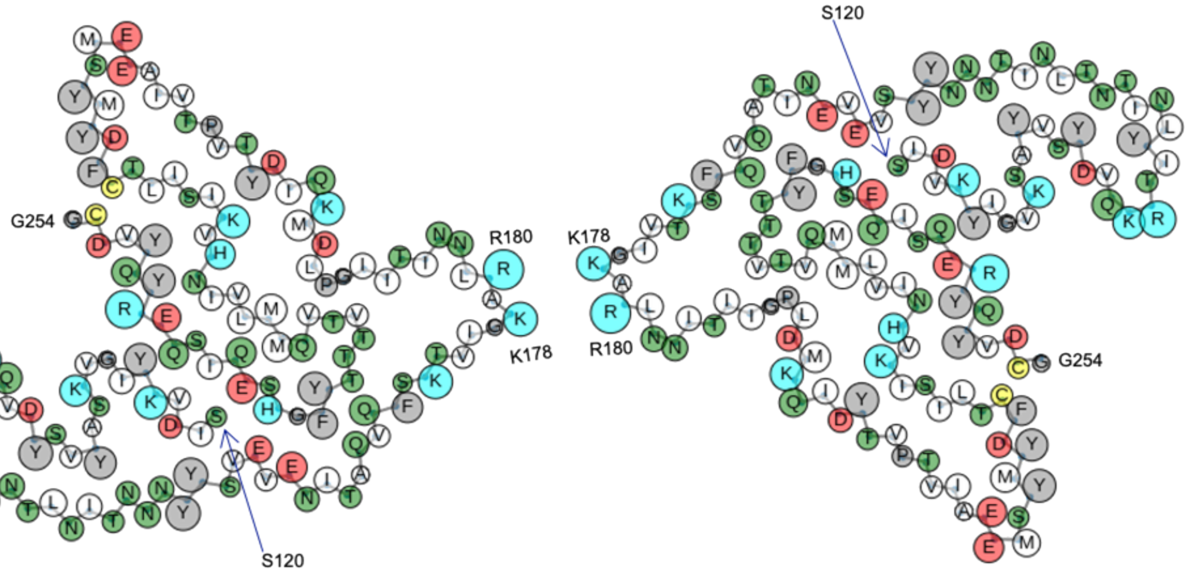


SFigure 4

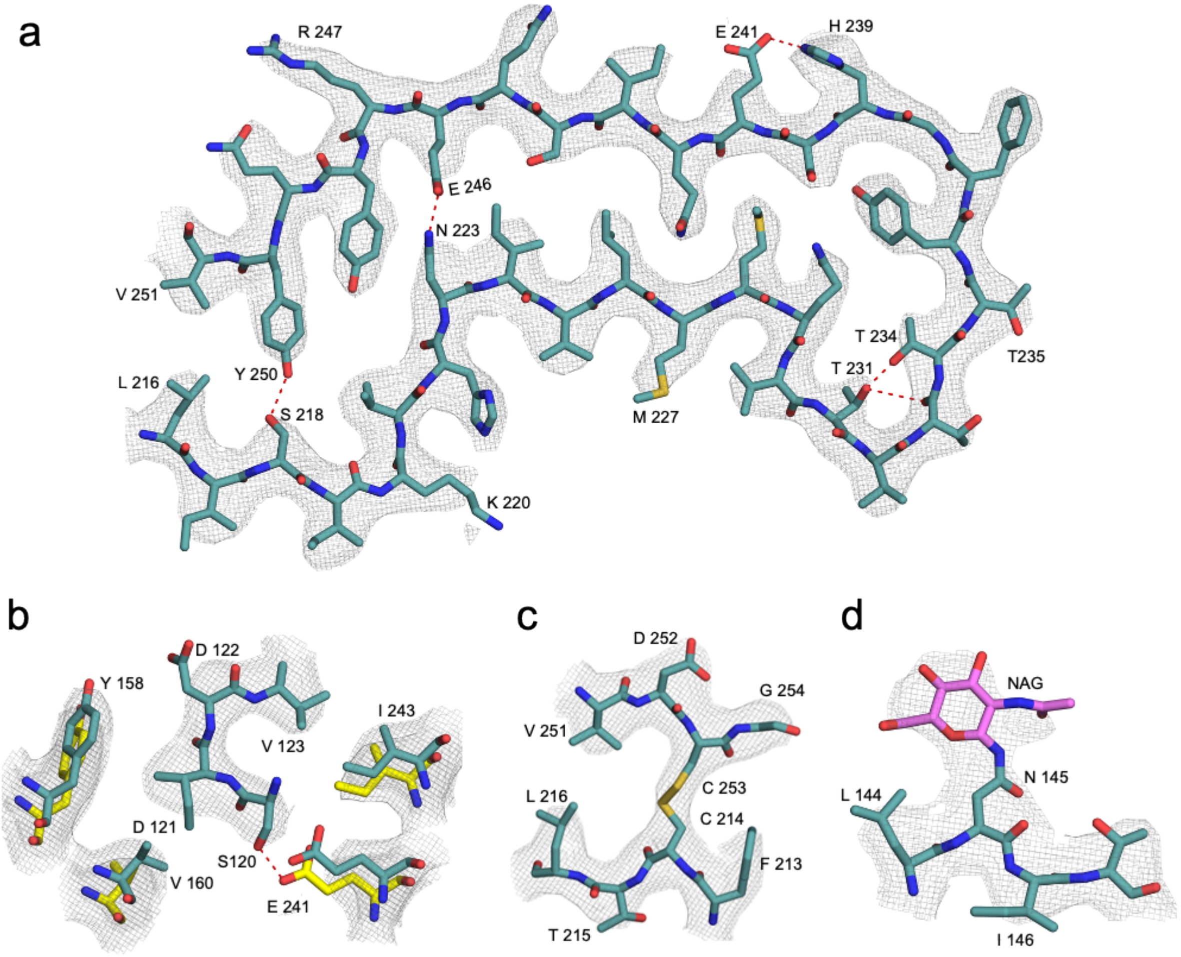
a
R 247
E 241
H 239
E 246
N 223
V 251
T 234
T 231
T235
Y 250
L 216
S 218
M 227
K 220
b
D 122
Y 158
V 123
I 243
D 121
S120
V 160
E 241
c
D 252
G 254
V 251
L 216
C 253
C 214
F 213
T 215
d
NAG
N 145
L 144
I 146

SFigure 5

**Supplementary Figures.**

**SFigure 1. Photomicrographs show neuronal and glial inclusions stained with Gallyas silver method and labeled with Tau antibodies.** Argyrophilic inclusions in neurons and glia of the gray matter (a,c). Argyrophilic inclusions in glia of the white matter (b,d). Inclusions in neurons and glia of the gray matter are labeled with anti-four-repeat tau (RD4) antibodies (e,g). Inclusions in glia of the white matter are labeled with RD4 antibodies (f,h). Neurons and glia of the gray matter are not labeled with anti-three-repeat tau (RD3) antibodies (i,k). Glial cells of the white matter are not labeled with RD3 antibodies (j,l). Gallyas silver method (a,b,c,d). Anti-tau antibody RD4 (e,f,g,h). Anti-tau antibody RD3 (i,j,k,l). Scale bar: 20 µm. Proteins identified by MS that co-purify with tau deposits from gray and white matter from cases #1 and #2 were analyzed. 2,006 proteins were present in all samples (m).

**SFigure 2. Cryo-EM imaging of tau and TMEM106B filaments in MSTD**. a) Representative cryo-EM images of filaments from sarkosyl-insoluble fractions from cases #1 and #2. Green arrows indicate tau filaments, red arrows indicate TMEM106B singlet filaments, and yellow arrows indicate TMEM106B doublets. b) Side-view of the cryo-EM 3D reconstructions. c) Power spectra of tau and TMEM106B singlet and doublets 2D class averages. Scale bars: a) 50 nm, b) 35 nm.

**SFigure 3. Fourier shell correlation curves and analyses using phenix.sequence_from_map for TMEM106B filaments.** a) Fourier shell correlation (FSC) curves for independently-refined maps for TMEM106B singlets (white matter cases #1 and #2, blue and yellow color respectively) and doublet (gray matter case #1, green). The FSC threshold of 0.143 is shown as the dash line. b) The LLG scores from phenix.sequence_from_map were plotted as a function of the sequence id and top-ranking sequence was identified as TMEM106B

**SFigure 4. Atomic model of TMEM106B.** The size of the circles are proportional to the radius of gyration computed for each residue. Large hydrophobic (gray), small hydrophobic (white), positively charged (teal), small polar (green), negatively charged (red), and cysteine (yellow) residues are highlighted.

**SFigure 5. Electron density of selected regions of TMEM106B.** The fibril structure is stabilized by the presence of hydrophobic zippers, hydrogen bonds between the layers and a disulphide bond. TMEM106B is glycated at four different positions. a) Zipper formed by two β-strands. b) Hydrogen bonds between two layers of TMEM106B. c) Disulfide bond between Cys 253 and Cys 214. d) Electron density of a glycated Asn145.

Table S1. Cryo-EM data collection, refinement and validation statistics.

| Data Collection | Gray matter case 1 | | | White matter case 1 | | | Gray matter case 2 | | | White matter case 2 | | |
|---|---|---|---|---|---|---|---|---|---|---|---|---|
| Magnification | x105000 | | | x105000 | | | x81000 | | | x81000 | | |
| Defocus range (µm) | -1.0 to -3.0 | | | -1.0 to -3.0 | | | -1.0 to -2.5 | | | -1.0 to -2.5 | | |
| Voltage (kV) | 300 | | | 300 | | | 300 | | | 300 | | |
| Microscope | Krios | | | Krios | | | Krios | | | Krios | | |
| Camera | K3 | | | K3 | | | K3 | | | K3 | | |
| Number movies collected | 11,145 | | | 11,980 | | | 23,345 | | | 8,336 | | |
| Total electron dose ($e^-/Å^2$) | 49.00 | | | 54.66 | | | 48.93 | | | 55.17 | | |
| Pixel size (Å) | 0.8354 | | | 0.8236 | | | 1.0554 | | | 1.0554 | | |
| **Reconstruction** | Singlet | Doublet | Tau | Singlet | Doublet | Tau | Singlet | Doublet | Tau | Singlet | Doublet | Tau |
| Box size (pixel) | 320 | 400 | 280 | 320 | 400 | 280 | 320 | 400 | 280 | 320 | 400 | 280 |
| Inter-box distance (Å) | 32 | 40 | 28 | 32 | 40 | 28 | 34 | 40 | 28 | 32 | 40 | 28 |
| Symmetry imposed | C1 | C2 | C2 | C1 | C2 | C2 | C1 | C2 | C2 | C1 | C2 | C2 |
| Helical rise (Å) | 4.79 | 4.79 | 4.79 | 4.78 | 4.79 | 4.79 | 4.81 | 4.79 | 4.79 | 4.79 | 4.80 | 4.78 |
| Helical twist (°) | -0.40 | -0.41 | -0.38 | -0.42 | -0.41 | -0.39 | -0.42 | -0.41 | -0.39 | -0.43 | -0.42 | -0.39 |
| Resolution (Å) | 8 | 7 | 6 | 3.6 | 3.8 | 4.6 | 3.3 | 3.9 | 5 | 2.90 | 3.4 | 4.0 |
| FSC threshold | 0.143 | 0.143 | 0.143 | 0.143 | 0.143 | 0.143 | 0.143 | 0.143 | 0.143 | 0.143 | 0.143 | 0.143 |

Table S2. Genetic variants identified by whole exome sequencing.

**Case 1**

| Gene | Variant | SNP ID |
|---|---|---|
| TMEM106B | S134N | RS147889591 |
| TMEM106C | S175F | RS2286025 |
| VPS11 | H34R | RS584115 |
| VPS11 | L35F | RS670526 |
| PLD3 | V323M | RS145999145 |
| APOE | N14K | RS440446 |
| APOE | R176C | RS7412 |
| KCNMA1 | E134Q | RS139968359 |
| KCNMA1 | S46delinsSS | RS572827902 |

**Case 2**

| Gene | Variant | SNP ID |
|---|---|---|
| TMEM106C | V103F | RS35000511 |
| VPS11 | K889R | RS15818 |
| VPS11 | L35F | RS670526 |
| APOE | N14K | RS440446 |
| KCNMA1 | S12G | RS77602559 |
